# Simultaneous knockout of VITAMIN C DEFECTIVE 2 and 3 exacerbates ascorbate deficiency and light stress sensitivity

**DOI:** 10.1101/2025.04.03.647133

**Authors:** Yasuhiro Tanaka, Takanori Maruta, Koki Arima, Hiroki Nakayama, Akane Hamada, Takahiro Ishikawa

## Abstract

Light-dependent activation of ascorbate biosynthesis is essential for accumulating ascorbate to mitigate photooxidative stress. The *VTC2* and *VTC3* genes play key roles in this process: *VTC2* encodes GDP-L-galactose phosphorylase (GGP), the rate-limiting enzyme in ascorbate biosynthesis, and *VTC3* encodes a putative protein kinase/phosphatase with an unknown function. Here, we investigated their functional and genetic relationship. In *vtc3* mutants, *VTC2* transcription and GGP activity were slightly enhanced, suggesting that VTC3 is not required for VTC2 expression. Additionally, the *vtc3* mutation had negligible effects on the transcriptome and the activity of enzymes involved in ascorbate redox cycle regulation, narrowing down the possible roles of VTC3 in ascorbate biosynthesis. Under low-light conditions, ascorbate levels were lower in *vtc2* than in *vtc3* mutants, but *vtc2* retained the ability to increase ascorbate levels under high-light stress, unlike *vtc3*. The simultaneous knockout of VTC2 and VTC3 further reduced ascorbate levels compared to single mutants and severely impaired light stress-induced ascorbate accumulation, resulting in impaired non-photochemical quenching and enhanced photooxidative damage. These findings highlight the additive effects of VTC2 and VTC3 on ascorbate biosynthesis and stress tolerance. The *vtc2 vtc3* double mutants provide a valuable model for further elucidating the physiological roles of ascorbate in plants.

## Introduction

In plants, reactive oxygen species (ROS) can be generated as byproducts of photosynthesis (Asada, 1999). Other major sources of ROS include the mitochondrial electron transport chain and the oxidation of glycolate in peroxisomes during photorespiration (Mittler *et al*., 2022). In addition, plants possess multiple apoplastic ROS-producing enzymes, particularly NADPH oxidases (also known as respiratory burst oxidase homologs) (Mittler *et al*., 2022). Under stress conditions, ROS generation through these various pathways is often inevitably increased or actively promoted as part of the plant’s response mechanisms. While ROS are highly reactive and can damage cellular components at elevated levels, they also serve as critical signaling molecules in plant stress acclimation (Mittler *et al*., 2022). Maintaining ROS homeostasis is essential for harnessing their signaling functions without compromising cellular integrity. This balance is achieved through robust antioxidant systems, among which ascorbate is one of the most abundant soluble antioxidants in plant leaves, present in all intracellular compartments at millimolar (mM) concentrations (Zechmann *et al*., 2011; Smirnoff, 2018; Maruta *et al*., 2024). Ascorbate plays a central role in scavenging ROS through both direct nonenzymatic reactions and enzymatic pathways mediated by ascorbate peroxidase (APX), an enzyme specific to eukaryotic photosynthetic organisms (Shigeoka *et al*., 2002; Maruta *et al*., 2016). Additionally, ascorbate serves as a substrate for violaxanthin de-epoxidase, an enzyme crucial for the xanthophyll cycle, which dissipates excess light energy as heat and limits ROS production during photosynthesis (Müller-Moulé *et al*., 2004). Beyond its role in ROS management, ascorbate is involved in diverse biological processes, including photosynthetic electron transport, iron uptake, cell division, anthocyanin accumulation, and hormone biosynthesis (Smirnoff, 2018).

Several pathways for ascorbate biosynthesis have been proposed; however, the D-mannose/L-galactose (D-Man/L-Gal) pathway, also known as the Smirnoff–Wheeler pathway, is widely recognized as the primary route (Wheeler *et al*., 1998; Smirnoff, 2018; Maruta, 2022). This notion is supported by substantial experimental evidence demonstrating its involvement in ascorbate biosynthesis across land plants and algae (Wheeler *et al*., 1998; Conklin *et al*., 1999; Vidal-Meireles *et al*., 2017; Sodeyama *et al*., 2021; Ishida *et al*., 2024). The D-Man/L-Gal pathway synthesizes ascorbate from the hexose phosphate pool through eight enzymatic reactions catalyzed sequentially by phosphomannose isomerase (PMI), phosphomannomutase (PMM), GDP-D-mannose pyrophosphorylase (GMP), GDP-D-mannose-3’,5’-epimerase (GME), GDP-L-galactose phosphorylase (GGP), L-galactose-1-phosphate phosphatase (GPP), L-galactose dehydrogenase (L-GalDH), and L-galactono-1,4-lactone dehydrogenase (L-GalLDH) (Smirnoff, 2018). The final step, catalyzed by L-GalLDH, occurs in the inner mitochondrial membrane, whereas all other enzymatic steps take place in the cytosol (Smirnoff, 2018). The significance of the D-Man/L-Gal pathway has been confirmed mainly through the characterization of Arabidopsis vitamin C-defective (*vtc*) mutants. The genes *VTC1*, *VTC2*, and *VTC4* encode GMP, GGP, and GPP, respectively (Conklin *et al*., 1999, 2006; Jander *et al*., 2002; Laing *et al*., 2007; Linster *et al*., 2007). Arabidopsis also possesses a minor isoform of GGP, encoded by *VTC5*. Notably, a double knockout mutant of *VTC2* and *VTC5* results in undetectable ascorbate levels and exhibits seedling lethality unless supplemented with ascorbate or its precursors (Dowdle *et al*., 2007).

Light is a potent stimulus for increasing the ascorbate pool size in plants. Leaf ascorbate content rises in response to light intensity and exposure duration (Dowdle *et al*., 2007; Yabuta *et al*., 2007). The key enzyme regulating ascorbate content under light conditions is VTC2/GGP, whose transcription is induced by light (Dowdle *et al*., 2007; Yabuta *et al*., 2007). Interestingly, the light-dependent activation of *VTC2* transcription is suppressed by photosynthesis inhibitors, suggesting a role for photosynthesis in the regulation of ascorbate biosynthesis (Yabuta *et al*., 2007). GGP activity is significantly enhanced under high-light stress, whereas the activities of other biosynthetic enzymes remain unaffected (Dowdle *et al*., 2007). Moreover, overexpression of VTC2 effectively enhances ascorbate pool sizes in various plant species (Bulley *et al*., 2009, 2012; Yoshimura *et al*., 2014). These findings confirm that VTC2/GGP is the rate-limiting enzyme in the D-Man/L-Gal pathway, a conclusion further supported by computational simulations (Fenech *et al*., 2021). The regulation of VTC2 by light also occurs at the post-translational level. In the dark, GGP activity is inhibited by a new class of blue-light receptors, PAS/LOV proteins, which directly associate with GGP (Ogura *et al*., 2008; Aarabi *et al*., 2023; Bournonville *et al*., 2023). Upon exposure to blue light, PAS/LOV proteins are dissociated from GGP, promoting its enzymatic activity. Thus, light-dependent activation of GGP plays a pivotal role in ascorbate biosynthesis. Recent studies have proposed that the biosynthetic enzymes in the cytosol form a protein complex (Fenech *et al*., 2021), potentially contributing to the regulation of ascorbate biosynthesis. Additionally, ascorbate itself tightly regulates its biosynthesis. High ascorbate levels repress VTC2 translation via an upstream open reading frame (uORF) in the 5′-untranslated region of the gene, counterbalancing the light-induced activation of GGP (Laing *et al*., 2015). Consequently, ascorbate biosynthesis is finely tuned through transcriptional, translational, and post-translational regulation. However, the precise mechanisms underlying light-dependent GGP activation remain to be elucidated.

The *VTC3* gene encodes a unique protein with both serine/threonine kinase and PP2C domains (Conklin *et al*., 2000, 2013). Two distinct *vtc3* alleles have been identified: *vtc3-1*, which carries a G202E missense mutation in the kinase domain, and *vtc3-2*, which has a nonsense mutation in the PP2C domain (Conklin *et al*., 2013). Both alleles exhibit a moderate ascorbate-deficient phenotype, with approximately 50% of wild-type ascorbate levels, a deficiency less severe than that observed in *vtc2* mutants (20–30% of wild-type levels). VTC3 is highly conserved across land plants and algae (Conklin *et al*., 2013; Maruta *et al*., 2024), and its loss also results in ascorbate deficiency in the moss *Physcomitrella patens* (Sodeyama *et al*., 2021). Notably, light-induced ascorbate accumulation is significantly impaired in Arabidopsis *vtc3* mutants (Conklin *et al*., 2013), suggesting that VTC3 plays a crucial role in the light regulation of VTC2/GGP. Since VTC3 has been reported to be a chloroplast protein, it is unlikely to regulate ascorbate biosynthesis through direct (de)phosphorylation of biosynthetic enzymes. The precise physiological function of VTC3 remains unclear, and further studies are required to elucidate its role.

The size of the ascorbate pool is regulated not only by biosynthesis but also by its oxidation, recycling, and degradation. Reduced ascorbate (ASC) undergoes oxidation through APX or direct interaction with ROS, generating monodehydroascorbate (MDHA) (Asada, 1999; Smirnoff, 2018). MDHA radicals are disproportionated into ASC and dehydroascorbate (DHA). MDHA and DHA are recycled back to ASC by monodehydroascorbate reductase (MDAR) and dehydroascorbate reductase (DHAR), respectively (Foyer and Halliwell, 1977; Hossain and Asada, 1985). MDAR uses NAD(P)H, while DHAR relies on reduced glutathione (GSH) as the electron donor. Additionally, GSH can nonenzymatically reduce DHA to ASC (Winkler *et al*., 1994; Terai *et al*., 2020; Hamada *et al*., 2023). However, DHA is highly unstable and undergoes irreversible degradation if not rapidly reduced. These interconnected processes are essential for maintaining ascorbate homeostasis. These interconnected processes are essential for maintaining ascorbate homeostasis, and it remains possible that VTC3 regulates ascorbate pool size by modulating ascorbate oxidation or degradation processes.

This study addresses two unresolved questions regarding the regulation of ascorbate homeostasis in plants. First, although VTC2/GGP is known to be a key regulatory enzyme in ascorbate biosynthesis and is strongly induced by light, the upstream signaling components controlling its activation remain poorly understood. Previous studies reported that *vtc3* mutants exhibit defective light-induced ascorbate accumulation (Conklin *et al*., 2013), suggesting a possible regulatory link between VTC3 and VTC2/GGP expression or activity. Based on this, we hypothesized that VTC3 facilitates the light-dependent activation of VTC2/GGP and thereby promotes ascorbate biosynthesis under light stress. Second, while *vtc2* and *vtc3* single mutants both show ascorbate deficiency, their distinct phenotypes raise the question of whether these genes act independently or synergistically. Notably, *vtc2* mutants retain the ability to increase ascorbate levels under high-light stress, likely due to the residual function of the minor GGP isoform VTC5 (Hamada *et al*., 2023; Terai *et al*., 2020), whereas *vtc3* mutants fail to induce this response (Conklin *et al*., 2013). We therefore hypothesized that the simultaneous loss of both genes would result in a more severe ascorbate-deficient phenotype and increased light stress sensitivity. By characterizing single and double mutants of *vtc2* and *vtc3*, we aimed to test these hypotheses and gain new insights into the genetic and physiological control of ascorbate metabolism and photoprotection in Arabidopsis.

## Materials and methods

### Plant Materials and Growth Conditions

*Arabidopsis thaliana* ecotype Col-0 was used as the wild type. The T-DNA insertion lines *vtc2-4* (SAIL_769_H05) and *vtc3-3* (SALK_095743) were obtained from the Arabidopsis Biological Resource Center and used for further crossings. Unless otherwise specified, wild-type and mutant seeds were sown in Jiffy-7 peat pellets and stratified in the dark at 4°C for 2–3 days. Plants were grown in a growth chamber (KCLP-1000CCFL-6-8L-CO₂, NK System) at 22°C under a 16-hour photoperiod with low light intensity (40–60 µmol photons m^-2^ s^-1^) and at 20°C in darkness for 8 hours. At three weeks of age, plants were exposed to moderate- or high-light stress (750 or 1,500 µmol photons m^-2^ s^-1^) using an LED lighting unit (PFQ-600PT, NK System) without a preceding dark period. The LED system minimized heat production, maintaining leaf temperature at approximately 24–25°C. To ensure uniform light exposure, plants were placed on a turntable beneath the LED panel. Fully expanded rosette leaves were collected from at least three plants and treated as a single biological replicate. Rosette and lesion areas (percentage of the total rosette area) were quantified using ImageJ software.

For ascorbate complementation assays, fully expanded leaves were excised from 3-week-old plants and floated on 20 mM MES buffer (pH 5.7), with or without 10 mM sodium ascorbate. The samples were incubated under continuous light for 24 h, followed by exposure to high-light conditions for an additional 24 h. Thereafter, leaves were subjected to trypan blue staining to visualize cell death. The stained area was quantified using ImageJ software.

For plate-grown experiments (**Figures 3 and 4**), wild-type and mutant plants were cultivated on half-strength Murashige and Skoog (MS) medium supplemented with 1% sucrose under a 16-hour photoperiod at 100 µmol photons m^-2^ s^-1^ and 22°C, with an 8-hour dark period at 20°C (LH-240S, NK System). Plants were then subjected to moderate-light stress (650 µmol photons m^-2^ s^-1^) without an intermediate dark period using the same LED panel and turntable system. Shoots from at least ten plants were pooled and treated as a single biological replicate.

### Semi-quantitative and quantitative reverse transcription-polymerase chain reaction

Semi-quantitative reverse transcription-polymerase chain reaction (RT-PCR) was performed according to Terai *et al*. (2020). Quantitative RT-PCR (qPCR) was performed as described by Iwagami *et al*. (2022). *Actin2* mRNA was used as an internal standard in all experiments. The expression stability of *Actin2* was confirmed by evaluating C*t* values across all experimental conditions. Three biological replicates per genotype/treatment combination were used, and each was derived from the average of three technical qPCR replicates. Expression variability was assessed using the coefficient of variation (CV). The consistently low CV values (ranging from 0.4% to 1.32%) support the use of *Actin2* as a stable reference gene under the tested conditions (see **Supplementary Figure S1**). All the primers used are listed in **Supplementary Table S1**.

### Western blotting

To examine VTC2 protein expression, we attempted to generate two independent anti-VTC2 peptide antibodies. The peptide sequences used were “NH_2_-C+QNSSGNVNQKSNRT-COOH” and “NH_2_-C+KRKEDYEGASEDNA-COOH”. The design and synthesis of the antigenic peptides, as well as antibody production, were outsourced to Eurofins Genomics. In addition, we purchased a commercial anti-GDP-L-galactose phosphorylase antibody (AS10 723, Agrisera). Using these antibodies, we performed western blotting following the method described by Kameoka *et al*. (2021), but we were unable to detect the VTC2 protein.

### Measurements of Antioxidants and Enzyme Activity

Ascorbate content was quantified using an ultra-fast liquid chromatography (UFLC) system equipped with a C18 column, following the method of Hamada and Maruta (2024). Glutathione levels were measured according to Noctor *et al*. (2016) without modifications. Soluble and membrane-bound APX activity was assessed using the method described by Kameoka *et al*. (2021), while DHAR and MDAR activities were measured as outlined by Tanaka *et al*. (2021).

GGP activity was measured according to Hamada *et al*., (2023). *Arabidopsis* leaves (0.2 g) frozen in liquid nitrogen were ground and homogenized in 1 mL of extraction buffer containing 50 mM Tris–HCl (pH 7.5), 10% glycerol, 2 mM DTT, 10 mM MgCl_2_, 1 mM aminocaproic acid, 1 mM benzamidine, 1 mM phenylmethanesulfonylfluoride. After centrifugation (15,300 *× g*) for 20 min at 4 °C, the supernatant was concentrated by ultrafiltration (15,300 *× g*, 30 min, 4 °C) using an Amicon ultra centrifugal filter (10 kDa, NMWL) and used for the following assay. The extract (20 µL) was added to 80 µL of a reaction mixture containing 50 mM Tris–HCl (pH 7.5), 2 mM MgCl_2_, 10 mM KCl, 1mM dithiothreitol, and 100 µM GDP-D-glucose with or without 5 mM sodium phosphate and incubated for 1 h at 26 °C. The reaction was stopped by boiling for 5 min, and the mixture was passed through 0.22 µm filters (Merck Millipore). The samples were injected into an ultra-fast liquid chromatography (UFLC) system (Prominence UFLC, Shimadzu) equipped with a C-18 column (LUNA C18(2), Column 150 x 4.6 nm, Shimadzu). The mobile phase consisted of 50 mM phosphate buffer (pH 6.7), 5 mM tetrabutylammonium dihydrogen phosphate, 5 mg L^−1^ EDTA, and 3.5% methanol, and the flow rate was 1 mL min^-1^. GDP-D-glucose was detected at 254 nm using the SPD-20A UV-Vis detector (Shimadzu). Pi-independent degradation was subtracted from the Pi-dependent decrease in the substrate to calculate GDP-D-glucose degradation activity.

### RNA-Seq Analysis

Two-week-old wild-type, *vtc2-4*, and *vtc3-3* seedlings, grown on half-strength MS medium supplemented with 1% sucrose under the same conditions described above, were incubated in darkness for two days, followed by a six-hour exposure to growth light (100 µmol photons m^-2^ s^-1^). Total RNA was extracted from shoot tissues using RNAiso Plus (Takara, Japan) and purified with NucleoSpin RNA Plus (Takara). RNA was fragmented and converted to double-stranded DNA using the TruSeq RNA Sample Preparation Kit v2 (Illumina). Library quality and quantity were assessed using a Bioanalyzer (Agilent) and a KAPA Library Quantification Kit (KAPA Biosystems), respectively. Sequencing was performed using an Illumina MiSeq with the MiSeq Reagent Kit v3 (300 cycles; Illumina). RNA-seq data were obtained from triplicate biological samples (n = 3 per group) and statistically analyzed. Read mapping, data processing, and statistical analyses were conducted using the CLC Genomics Workbench (version 9.5.2). The gene expression dataset has been deposited in the Gene Expression Omnibus under accession number GSE293768. Gene ontology (GO) enrichment analysis was performed using agriGO (version 2; Tian *et al*., 2017).

### Chlorophyll Fluorescence Measurements

Chlorophyll fluorescence measurements were performed using a Closed FluorCam 800MF (Photon Systems Instruments). Plants were dark-adapted for 30 minutes before measurements. Chlorophyll fluorescence parameters, F_v_/F_m_ and NPQ, were calculated according to Baker (2008) using the following equations: F_v_/F_m_ = (F_m_ - F_o_)/F_m_, and NPQ = (F_m_ – F_m_ʹ)/F_m_ʹ. NPQ was measured under actinic light at an intensity of 90 µmol photons m^-2^ s^-1^.

### Accession Numbers

VTC2, At4g26850; VTC3, At2g40860.

### Data Analysis

The statistical methods used are described in the respective figure legends (Tukey–Kramer, Dunnett’s test, or Student’s *t*-test). Statistical calculations were performed using at least three independent biological replicates (see figure legends for details).

## Results

### VTC3 is required for ascorbate accumulation under high-light stress

To evaluate the role of VTC3 in ascorbate accumulation under high-light conditions, we used a T-DNA insertion mutant for the *VTC3* gene, referred to hereafter as *vtc3-3* (Conklin *et al*., 2013), as well as the *vtc2-4* mutant (Lim *et al*., 2016). In *vtc3-3*, the T-DNA is inserted in the seventh exon of *VTC3*, whereas in *vtc2-4*, the insertion occurs in the third exon of *VTC2*. Both mutants have been previously characterized as null alleles, lacking full-length transcripts of the respective genes (Conklin *et al*., 2013; Lim *et al*., 2016). This was further confirmed by our RT-PCR analysis (**Supplementary Figure S2**). Under low-light conditions, *vtc2-4* and *vtc3-3* mutants exhibited approximately 26% and 45% of wild-type ascorbate levels, respectively (**Figure 1A**), consistent with previous reports (Conklin *et al*., 2013; Lim *et al*., 2016). Exposure to high-light stress induced a time-dependent increase in ascorbate content in wild-type plants. In contrast, the *vtc3-3* mutant displayed only a minimal and statistically insignificant increase in ascorbate levels. The ascorbate pool size in *vtc2-4* remained significantly lower than in wild-type plants under both low-light and high-light conditions. However, high-light stress markedly increased ascorbate levels in *vtc2-4* (**Figure 1A**), corroborating previous findings (Hamada *et al*., 2023; Terai *et al*., 2020). The elevated amounts of ascorbate after 48 hours of high-light exposure were approximately 5.5 µmol g^-1^ FW in the wild type, 3.4 µmol g^-1^ FW in *vtc2-4*, and 0.7 µmol g^-1^ FW in *vtc3-3* (**Figure 1B**). Neither the *vtc2-4* nor *vtc3-3* mutation affected the ascorbate redox state under either low- or high-light conditions. These results demonstrate that VTC3 is essential for the accumulation of ascorbate in response to high-light exposure.

**Figure 1.**
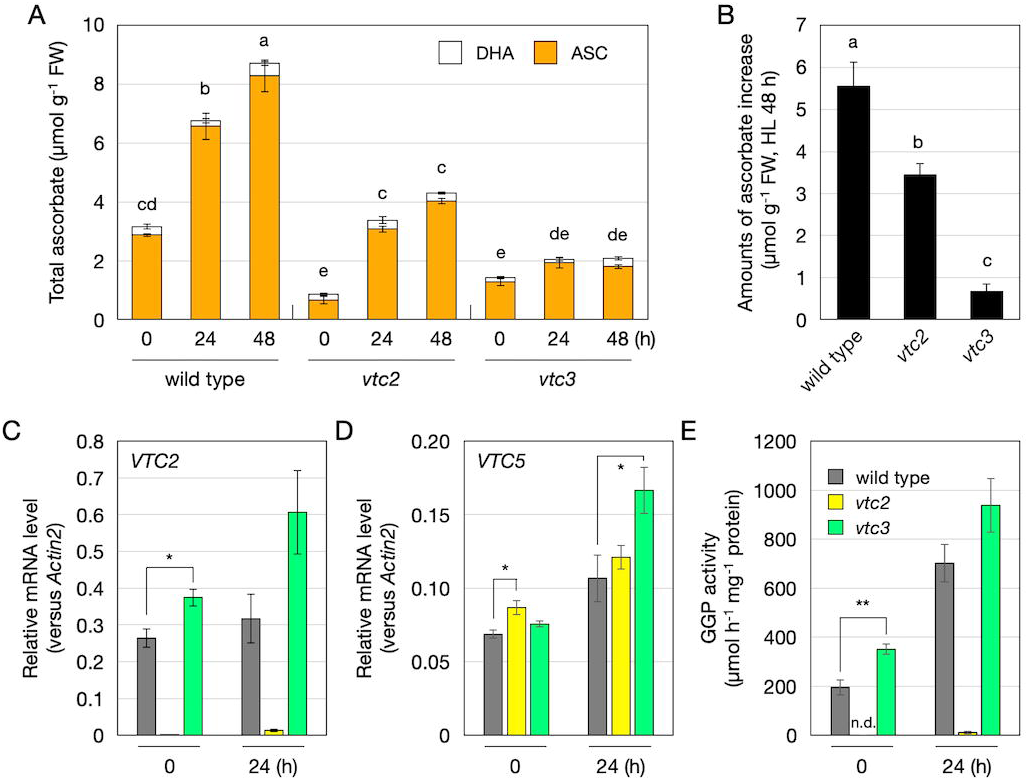
Ascorbate accumulation and GGP expression/activity in *vtc3-3* under high light Wild-type, *vtc2-4*, and *vtc3-3* plants were grown in soil under low-light (40–60 μmol photons m^−2^ s^−1^) conditions for three weeks and then exposed to high-light stress (1,500 μmol photons m^−2^ s^−1^) for 48 h. Fully expanded leaves were used in subsequent experiments. (A) Total ascorbate content (sum of ASC and DHA contents). (B) The amounts of ascorbate increase from before to after high-light exposure (48 h). (A, B) Data are presented as the mean ± SE of four biological replicates. Different letters indicate significant differences (*P* < 0.05, one- or two-way ANOVA followed by Tukey–Kramer test). (C, D) Transcript levels of *VTC2* and *VTC5* were determined using qPCR analysis. (E) GGP activity was determined as GDP-D-glucose degrading activity. Data are presented as the mean ± SE of three biological replicates. (C, E) Significant differences between wild type and *vtc3-3* (Student’s *t*-test): \**P* < 0.05, \*\**P* < 0.01, versus the values of the wild-type plants. Because *VTC2* transcript levels and GGP activity were negligible in *vtc2-4*, statistical analysis was performed only between wild-type and *vtc3-3*. (B) Different letters indicate significant differences between genotypes (*P* < 0.05, one-way ANOVA followed by Tukey–Kramer test). ASC, reduced ascorbate; DHA, dehydroascorbate; FW, fresh weight.

### VTC3 is not required for VTC2 expression or GGP enzyme activity

To test our first hypothesis that VTC3 regulates the light-dependent activation of VTC2/GGP, we analyzed the transcript levels of *VTC2* and *VTC5* before and after 24 hours of high-light exposure. In *vtc2-4* mutants, *VTC2* transcripts were nearly undetectable (**Figure 1C**). In wild-type plants, the transcript levels of both *VTC2* and *VTC5* showed a modest increase in response to high-light stress. Intriguingly, these increases were more pronounced in *vtc3-3* mutants. This upregulation of *VTC2* and *VTC5* in *vtc3-3* mutants may represent a compensatory response to ascorbate deficiency. However, no similar increase in *VTC5* expression was observed in *vtc2-4* mutants after high-light exposure (**Figure 1D**), possibly because *vtc2-4* mutants retained higher ascorbate levels than *vtc3-3* mutants under high-light conditions (**Figure 1A**).

To assess VTC2 protein levels, we obtained three anti-VTC2 antibodies targeting different VTC2 peptides. Unfortunately, none of these antibodies successfully detected VTC2 via western blotting, likely due to low antibody specificity or the very low abundance of VTC2 protein. As an alternative, we measured GGP activity using a Pi-dependent GDP-D-glucose degradation assay, a reliable indicator of endogenous GGP activity (Hamada *et al*., 2023). In wild-type plants, GGP activity increased in response to high-light exposure, whereas in *vtc2-4* mutants, activity was nearly undetectable (**Figure 1E**). Notably, the absence of VTC3 did not reduce GGP activity; rather, it appeared to enhance it, consistent with the observed increase in *VTC2* and *VTC5* transcript levels in *vtc3-3* mutants. Contrary to our hypothesis, these results indicate that VTC3 is not essential for the expression and activity of GGP.

### VTC3 does not affect ascorbate redox enzyme activities or transcriptome profiles

We next speculated that VTC3 may contribute to ascorbate redox cycling rather than biosynthesis. To test this possibility, we examined the impact of VTC3 on glutathione levels and the activities of APX, DHAR, and MDAR. Plants grown under plate conditions were used for these experiments. Two-week-old wild-type, *vtc2-4*, and *vtc3-3* plants on agar plates were exposed to moderate-light stress (650 µmol photons m^-2^ s^-1^; ML^650^) for 24 hours. This stress increased ascorbate levels in both wild-type and *vtc2-4* plants, but this response was nearly absent in *vtc3-3* mutants, consistent with the data obtained from *in vivo* soil experiments (**Figure 2**).

**Figure 2.**
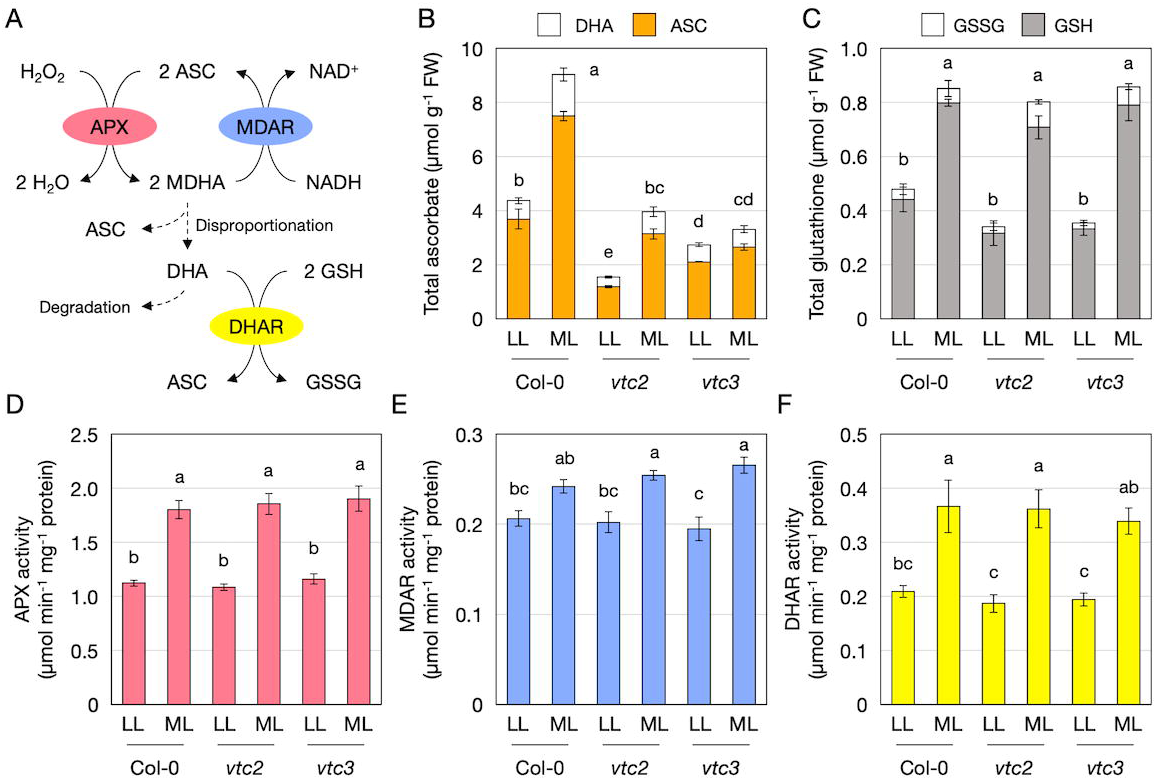
Antioxidant levels and enzyme activities related to ascorbate redox turnover in *vtc3-3* under high light Wild-type, *vtc2-4*, and *vtc3-3* plants were grown on MS plates under low-light (40–60 μmol photons m^−2^ s^−1^) conditions for two weeks and then exposed to moderate-light stress (650 μmol photons m^−2^ s^− 1^) for 24 h. (A) Ascorbate oxidation by ascorbate peroxidase (APX) and its recycling through monodehydroascorbate reductase (MDAR) and dehydroascorbate reductase (DHAR). (B) Total ascorbate content (sum of ASC and DHA contents). (C) Total glutathione content (sum of GSH and GSSG contents). (D) APX activity. (E) MDAR activity. (F) DHAR activity. Data are presented as the mean ± SE of three or four biological replicates. Different letters indicate significant differences (*P* < 0.05, two-way ANOVA followed by Tukey–Kramer test). ASC, reduced ascorbate; DHA, dehydroascorbate; FW, fresh weight; GSH, reduced glutathione; GSSG, oxidized glutathione.

The size of the glutathione pool increased under ML^650^ stress in all genotypes, with no significant differences observed among the wild type, *vtc2-4*, and *vtc3-3*. Additionally, APX, DHAR, and MDAR activities were upregulated under these conditions, but neither *vtc2-4* nor *vtc3-3* mutations influenced their activities under either low-light or stress conditions (**Figure 2**). These findings suggest that VTC3 does not significantly affect ascorbate redox cycling.

To further investigate the function of VTC3, we performed transcriptome analysis. For this experiment, a drastic light/dark shift was used instead of light stress to induce a stronger response. Two-week-old wild-type, *vtc2-4*, and *vtc3-3* plants grown on agar plates under low-light conditions were incubated in darkness for two days, followed by a 6-hour exposure to growth light (100 µmol photons m^-2^ s^-1^). Differentially expressed genes (false discovery rate < 0.05) were identified from triplicate datasets by comparing the mutants to wild-type plants (**Figure 3**). In total, over 200 genes were differentially expressed in *vtc2-4* mutants, whereas only 24 genes, including *VTC3* itself, were affected in *vtc3-3* mutants. Among these, seven genes were commonly affected in both mutants. The relatively minor transcriptomic impact of VTC3 compared to VTC2 likely reflects the weaker ascorbate-deficient phenotype of *vtc3-3*. Gene ontology (GO) enrichment analysis showed that “defense response to pathogens” was the most enriched GO term among the 12 genes upregulated in *vtc3* mutants (**Figure 3**). Furthermore, downregulated genes specific to *vtc3-3* included cytochrome P450 superfamily protein (AT3G44970), transmembrane protein (AT5G01881), mto1 responding down 1 (AT1G53480), and phosphoinositide 4-kinase gamma 1 (AT2G40850), none of which are known to be directly associated with ascorbate metabolism. These results suggest that VTC3 plays a more limited role in transcriptomic regulation than VTC2.

**Figure 3.**
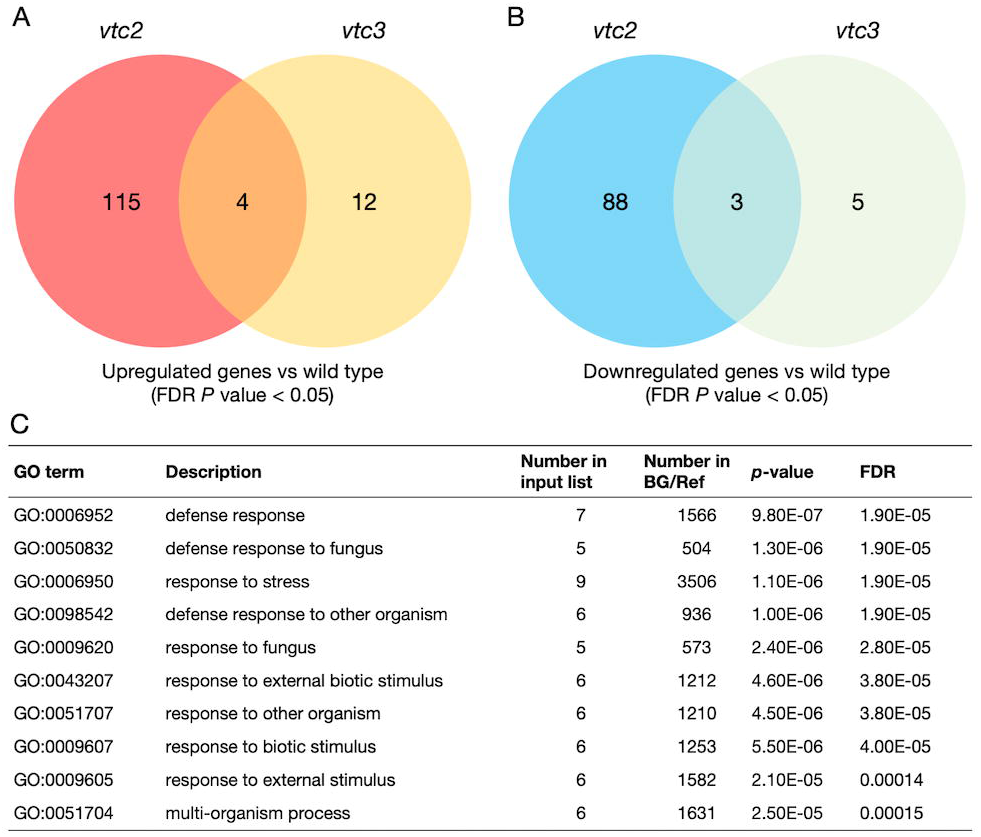
Transcriptomic changes in *vtc2-4* and *vtc3-3* Wild-type, *vtc2-4*, and *vtc3-3* plants were grown on MS plates under low-light (40–60 μmol photons m^−2^ s^−1^) conditions for two weeks and then incubated in darkness for two days (to decline ascorbate pool size), followed by a 6-hour exposure to light (100 µmol photons m^-2^ s^-1^). From the triplicate data, we selected the genes whose expression was significantly up-regulated (false discovery rate < 0.05) in *vtc2-4* or *vtc3-3* compared to the wild-type plants. (A, B) The Venn diagrams show the overlap between sets of genes upregulated (A) or downregulated (B) in *vtc2-4* and *vtc3-3*. (C) Gene ontology enrichment analysis (biological process) of 12 genes whose expression was upregulated in *vtc3-3* compared to the wild-type plants. (C) Gene ontology (GO) terms enriched in genes whose expression was specifically promoted in *vtc3-3*. BG, background; Ref, reference.

### Simultaneous loss of VTC2 and VTC3 exacerbates ascorbate deficiency and increases light stress sensitivity

To test our second hypothesis that the simultaneous loss of VTC2 and VTC3 would result in additive or synergistic effects on ascorbate pool size, we generated and analyzed *vtc2-4 vtc3-3* double mutants under both low- and high-light conditions. Homozygous double mutants were identified by genotyping the F_2_ population following crossing (**Supplementary Figure S2**). Compared to single mutants, the double mutants exhibited the lowest ascorbate levels, retaining only approximately 51% of the ascorbate content observed in *vtc2-4* single mutants, equivalent to about 14% of wild-type levels (**Figure 4**). Notably, except for mutants entirely lacking detectable ascorbate (e.g., *vtc2 vtc5* double mutants), *vtc2-4 vtc3-3* double mutants displayed the lowest ascorbate levels reported among ascorbate-deficient mutants. Despite these extremely low levels, their growth under low-light conditions for three weeks was comparable to that of wild-type plants and single mutants (**Figure 4**), suggesting that even 14% of wild-type ascorbate levels is sufficient to support Arabidopsis growth under controlled laboratory conditions.

**Figure 4.**
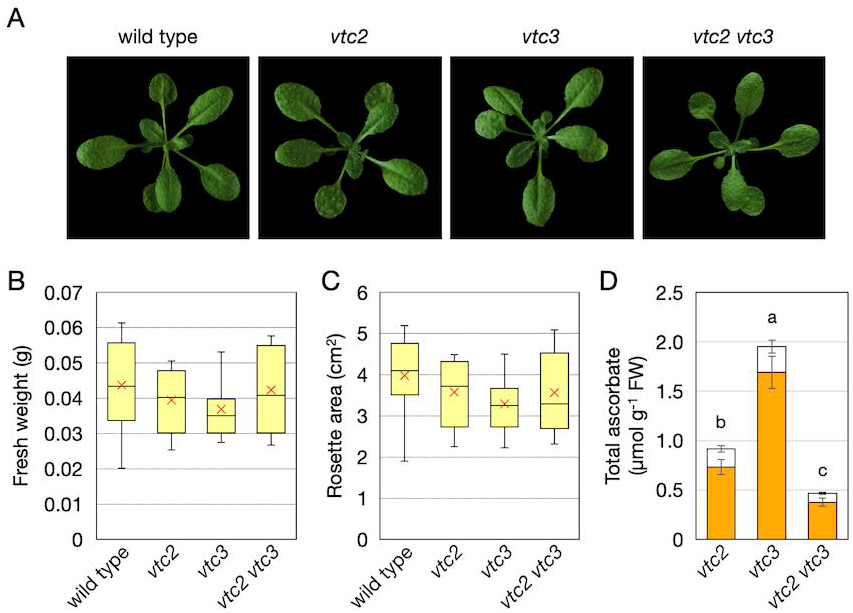
Phenotypes and ascorbate profiles of *vtc2-4 vtc3-3* double mutants Wild-type, *vtc2-4*, *vtc3-3*, and *vtc2-4 vtc3-3* plants were grown in soil under low-light (40–60 μmol photons m ^−2^ s ^−1^) conditions for three weeks. Fully expanded leaves were used for ascorbate measurements. (A) Representative phenotypes of plants. (B, C) Box plots illustrate fresh weights (B) and the rosette area (C). Horizontal bars and cross marks within the box represent median and mean values from 12 biological replicates, respectively, with boxes representing the 25th and 75th percentiles. The outer error bars (whiskers) represent maximum and minimum values. There was no significant difference between genotypes (one-way ANOVA). (D) Total ascorbate content (sum of ASC and DHA contents). Data are presented as the mean ± SE of five biological replicates. Different letters indicate significant differences (*P* < 0.05, one-way ANOVA followed by Tukey–Kramer test). ASC, reduced ascorbate; DHA, dehydroascorbate; FW, fresh weight.

When exposed to high-light stress for 24 hours, *vtc2-4* and *vtc3-3* single mutants exhibited photo-bleaching, primarily in older leaves (**Figure 5**). In contrast, the *vtc2-4 vtc3-3* double mutants displayed a more severe photo-bleaching phenotype, extending to mature leaves, and exhibited the most pronounced cell death, consistent with their extremely low ascorbate content. Reflecting these phenotypes, the reduction in the maximum quantum yield of photosystem II (F_v_/F_m_) under high-light stress was most pronounced in the double mutants (**Figure 5**), indicating severe damage to photosynthesis. To test whether the heightened light sensitivity of the double mutants is attributable to ascorbate deficiency, we conducted a chemical complementation experiment using sodium ascorbate. Detached leaves from 3-week-old *vtc2-4 vtc3-3* plants were floated on buffer with or without 10 mM sodium ascorbate under continuous light for 24 h, followed by 24 h of high-light exposure. Under these conditions, clear leaf bleaching and intense trypan blue staining were observed only in *vtc2-4 vtc3-3* leaves without sodium ascorbate treatment. Notably, these symptoms were completely abolished by sodium ascorbate treatment (**Supplementary Figure S3**), indicating that the severe photooxidative damage in the double mutants is primarily due to their profound ascorbate deficiency.

**Figure 5.**
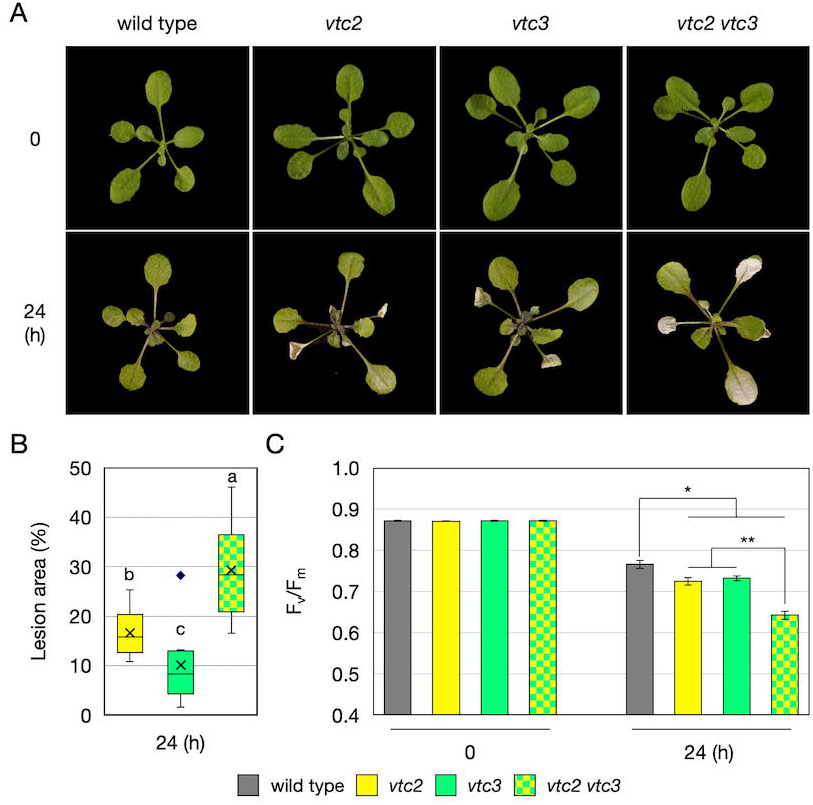
Sensitivity of *vtc2-4 vtc3-3* double mutants to high-light stress Wild-type, *vtc2-4*, *vtc3-3*, and *vtc2-4 vtc3-3* plants were grown in soil under low-light (40–60 μmol photons m^−2^ s^−1^) conditions for three weeks and then exposed to high-light stress (1,500 μmol photons m^−2^ s^−1^) for 24 h. (A) Representative phenotypes of plants before and after high-light stress. (B) Box plots illustrate lesion area (% of whole rosette area). The data were obtained from nine plants per genotype in three independent experiments. As wild-type plants showed no lesions, lesion area analysis was not performed. Horizontal bars and cross marks within the box represent median and mean values from five biological replicates, respectively, with boxes representing the 25^th^ and 75^th^ percentiles. The outer error bars (whiskers) and diamonds represent the maximum and minimum values, and an outlier, respectively. Different letters indicate significant differences (*P* < 0.05, one-way ANOVA followed by Tukey–Kramer test). (C) The maximum quantum yield of photosystem II (F_v_/F_m_) before and after high-light stress. Significant differences between genotypes (one-way ANOVA followed by Tukey–Kramer test): \**P* < 0.05, \*\**P* < 0.01.

Considering the heightened sensitivity of *vtc2-4 vtc3-3* mutants to high-light stress, we measured ascorbate content in wild-type plants and mutants during the early stages of high-light exposure (6 and 12 hours). Ascorbate levels in wild-type plants and *vtc2-4* mutants increased approximately 1.5-fold by 12 hours after high-light exposure (**Figure 6**), but there was no statistical difference. Furthermore, no significant increase in ascorbate levels was observed in *vtc3-3* or *vtc2-4 vtc3-3* mutants. However, the early-stage ascorbate increase in wild-type plants and *vtc2-4* mutants was relatively modest (**Figure 6**), making detailed comparisons between mutants challenging.

**Figure 6.**
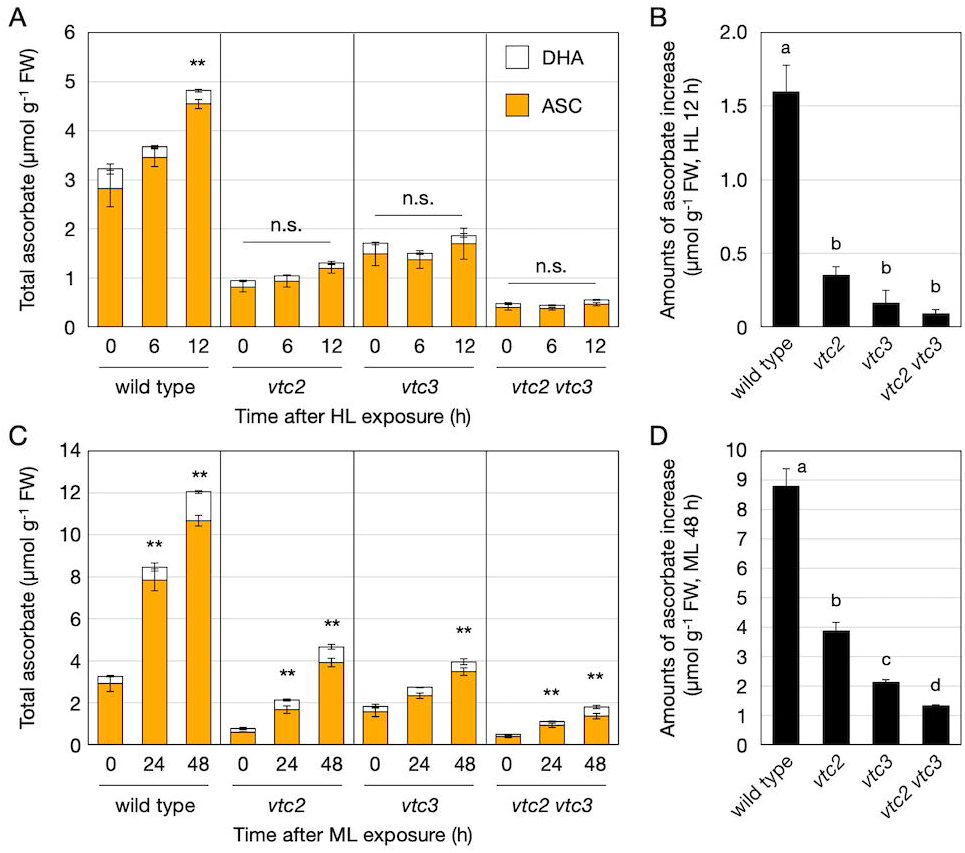
Ascorbate profiles in *vtc2-4 vtc3-3* under high- or moderate-light stress Wild-type, *vtc2-4*, *vtc3-3*, and *vtc2-4 vtc3-3* plants were grown in soil under low-light (40–60 μmol photons m^−2^ s^−1^) conditions for three weeks and then exposed to high-light stress (1,500 μmol photons m^−2^ s^−1^) for 24 h (A, B), or to moderate-light stress (750 μmol photons m^−2^ s^−1^) for 48 h (C, D). Fully expanded leaves were used for ascorbate measurements. (A, C) Total ascorbate content (sum of ASC and DHA contents). Data are presented as the mean ± SE of three or four biological replicates. Significant differences between data before and after light stress for each genotype (one-way-ANOVA with Dunnett’s post hoc test for total content): *P*\* < 0.05, \*\**P* < 0.01, compared to the values of 0 h. (B, D) The amounts of ascorbate increase from before to after high- or moderate-light exposure (24 or 48 h, respectively). Data are presented as the mean ± SE of three or four biological replicates. Different letters indicate significant differences (*P* < 0.05, one-way ANOVA followed by Tukey–Kramer test). ASC, reduced ascorbate; DHA, dehydroascorbate; FW, fresh weight; HL, high light; ML, moderate light; n.s., not significant.

To further investigate the ascorbate accumulation response under different light intensities, we conducted similar experiments under moderate-light conditions (750 µmol photons m^-2^ s^-1^; ML^750^). Under ML^750^ stress, ascorbate accumulation was observed even in *vtc3-3* and *vtc2-4 vtc3-3* mutants; however, the increase was significantly smaller compared to *vtc2-4*, with the double mutants exhibiting the lowest levels of stress-induced ascorbate accumulation (**Figure 6**). These results demonstrate that the simultaneous knockout of VTC2 and VTC3 severely limits both steady-state ascorbate levels and stress-induced ascorbate accumulation, thereby dramatically increasing sensitivity to photooxidative stress.

### Impaired induction of non-photochemical quenching in *vtc2-4 vtc3-3* double mutants

As ascorbate is implicated in the induction of NPQ (Müller-Moulé *et al*., 2004), a photoprotective mechanism that dissipates excess absorbed light energy as heat, we investigated how the severe ascorbate deficiency in *vtc2-4 vtc3-3* double mutants affects NPQ. Plants were grown under low-light conditions for three weeks before being subjected to high-light stress for 24 hours. Under low-light conditions, NPQ induction was partially suppressed in all mutants compared to wild-type plants, with statistically significant but relatively small differences (**Figure 7A and C**). Among the single mutants, *vtc2-4* exhibited a greater reduction in NPQ induction compared to *vtc3-3*, consistent with its lower ascorbate pool size. Although the *vtc2-4 vtc3-3* double mutants showed a trend toward further reduced NPQ values, the difference was not statistically significant compared to *vtc2-4* single mutants. Following high-light exposure, NPQ suppression was most pronounced in *vtc2-4 vtc3-3* double mutants, significantly lower than in either *vtc2-4* or *vtc3-3* single mutants (**Figure 7B and D**). Interestingly, while *vtc2-4* exhibited lower NPQ values than *vtc3-3* under low-light conditions (**Figure 7C**), this difference disappeared under high-light stress. (**Figure 7D**) These variations in NPQ induction closely paralleled the differences in ascorbate pool sizes among the mutants. These findings provide new insights into the relationship between NPQ induction and ascorbate levels, highlighting the additive effects of *vtc2-4* and *vtc3-3* mutations on NPQ suppression under high-light stress. The results suggest that ascorbate plays a critical role in NPQ induction under both low- and high-light conditions and that VTC2 and VTC3 contribute to this regulation through their roles in ascorbate accumulation.

**Figure 7.**
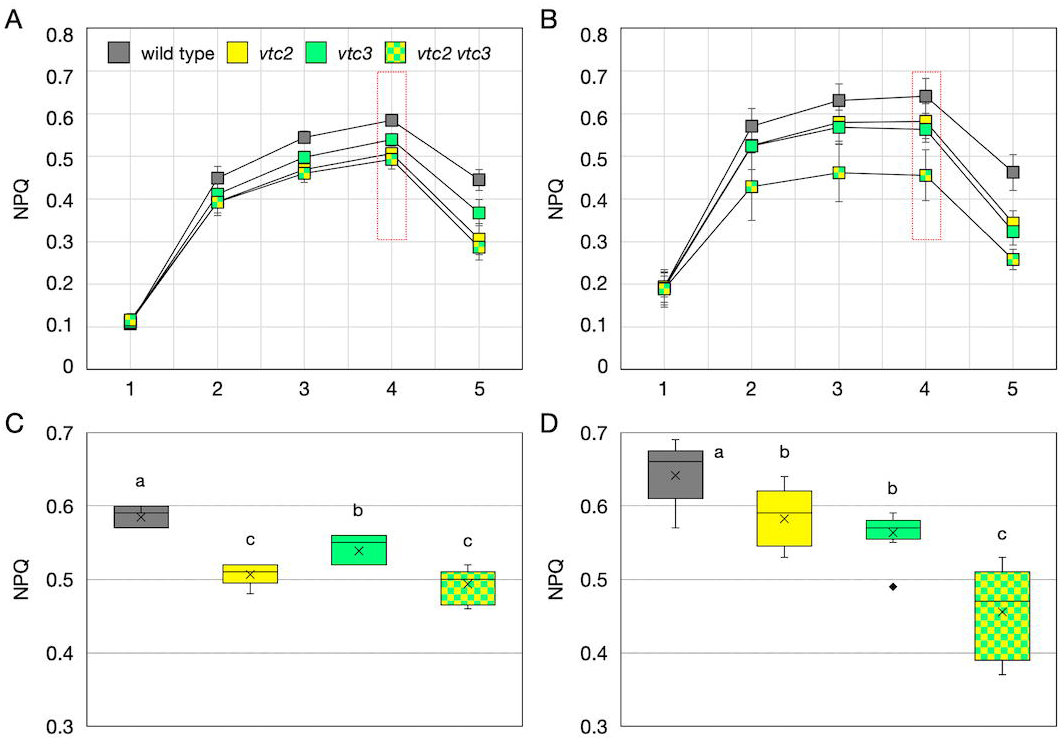
Non-photochemical quenching in *vtc2-4 vtc3-3* before and after high-light stress Wild-type, *vtc2-4*, *vtc3-3*, and *vtc2-4 vtc3-3* plants were grown in soil under low-light (40–60 μmol photons m^−2^ s^−1^) conditions for three weeks and then exposed to high-light stress (1,500 μmol photons m^−2^ s^−1^) for 24 h. (A, B) Non-photochemical quenching (NPQ) induction upon the onset of actinic light (90 µmol photons m^-2^ s^-1^) before (A) and after (B) stress. Data are presented as the mean ± SE of three or four biological replicates. The red triangles indicate the data points used for statistical analysis (C, D). (C, D) Box plots illustrate NPQ data before (C) and after (D) stress, corresponding to the areas outlined with red dashed lines in (A) and (B), respectively. Horizontal bars and cross marks within the box represent median and mean values from 12 biological replicates, respectively, with boxes representing the 25th and 75th percentiles. The outer error bars (whiskers) represent maximum and minimum values. Different letters indicate significant differences (*P* < 0.05, one-way ANOVA followed by Tukey–Kramer test).

## Discussion

Among the *vtc* mutant series, *vtc3* is the only mutant with a mutation in a gene that does not encode a biosynthetic enzyme (Conklin *et al*., 2013). VTC3 is a unique protein containing both protein kinase and phosphatase domains and is essential for ascorbate accumulation in response to continuous exposure to growth light. Given that VTC2/GGP is the rate-limiting enzyme in ascorbate biosynthesis and that its expression and activity are enhanced by light, we initially hypothesized that VTC3 regulates ascorbate biosynthesis at the VTC2/GGP step. However, this hypothesis was strongly refuted, as the loss of VTC3 did not reduce *VTC2/VTC5* transcription or GGP activity (**Figure 1**). As previously reported, VTC2 translation is controlled by ascorbate via an uORF in the 5′-untranslated region of the *VTC2* gene (Laing *et al*., 2015). Because we were unable to examine VTC2/GGP expression at the protein level, we cannot entirely rule out the possibility that VTC3 influences VTC2 translation. However, if this were the case, the absence of VTC3 would likely lead to reduced GGP activity, which was not observed (**Figure 1**). Thus, we conclude that VTC3 is not required for the expression and activity of VTC2/GGP.

We also found that the loss of VTC3 did not affect glutathione profiles or the activities of APX, DHAR, and MDAR (**Figure 2**). Since we did not examine the expression or activity of individual isoforms of these enzymes, we cannot rule out the possibility that VTC3 affects ascorbate redox cycling in specific compartments. However, recent studies have demonstrated that the complementary actions of MDARs, DHARs, and glutathione maintain a robust ascorbate recycling system in Arabidopsis (Maruta *et al.,* 2024). For instance, the loss of all three *DHAR* genes has a negligible impact on ascorbate levels, even under stress conditions (Rahantaniaina *et al*., 2017; Terai *et al*., 2020). Considering the resilience of the ascorbate recycling system, it is unlikely that VTC3 plays a role in regulating ascorbate redox cycling. Another possible function of VTC3 could be the inhibition of ascorbate degradation. However, this is also unlikely, as ascorbate degradation begins with DHA, and ascorbate recycling capacity was unaffected in *vtc3* mutants.

Since VTC3 contains both protein kinase and phosphatase domains, it was also plausible that it mediates light-dependent signaling pathways to enhance ascorbate pool size. However, transcriptome analysis revealed that only a limited number of genes were specifically up- or downregulated in *vtc3-3* mutants (**Figure 3**), and none had known associations with ascorbate metabolism. However, one of the genes downregulated in *vtc3-3* mutants encodes phosphoinositide 4-kinase gamma 1 (PI4Kγ1). Although this gene has not been directly linked to ascorbate metabolism, it may affect the pool size of *myo*-inositol, a potential precursor of ascorbate in the alternative biosynthetic pathway (Lorence et al., 2004). While the physiological relevance of this pathway in Arabidopsis remains under debate (Maruta, 2022), the potential role of PI4Kγ1 in regulating ascorbate levels through myo-inositol metabolism warrants further investigation. GO enrichment analysis identified “defense response to pathogens” as the most enriched term among the 12 genes whose expression was specifically enhanced in *vtc3* mutants (**Figure 3 and Supplementary Table S2**). This finding aligns with previous observations in *vtc1* mutants, where ascorbate deficiency was linked to enhanced pathogen responses (Pastori *et al*., 2003). Thus, the transcriptional activation of some defense-related genes in *vtc3-3* mutants is likely a secondary effect of ascorbate deficiency rather than the direct involvement of VTC3 in defense signaling. In *vtc2-4* mutants, 115 genes were upregulated (**Figure 3**), with GO terms related to “response to salicylic acid” and “response to chitin” enriched (**Supplementary Table S2**), further suggesting a link between ascorbate deficiency and pathogen responses.

In our RNA-seq analysis, we identified 210 differentially expressed genes (DEGs) in the *vtc2-4* mutant (**Figure 3**). This number is substantially lower than the 963 DEGs previously reported in *vtc2* mutants by Kerchev *et al*. (2011), who used microarray analysis. This discrepancy is likely due to differences in experimental conditions. In our study, 2-week-old seedlings were grown aseptically on agar plates under long-day conditions, followed by 48 hours of darkness and a 6-hour light treatment before sampling. In contrast, Kerchev *et al*. (2011) used fully mature 6-week-old plants grown in soil under short-day conditions. These differences in plant age, growth medium, and light regime likely contributed to the differing transcriptome responses. Moreover, only 11 out of the 119 upregulated genes in *vtc2-4* overlapped with the 378 upregulated genes in the *vtc2* dataset from Kerchev *et al*. (2011) (**Supplementary Table S2**), suggesting the condition-specific nature of transcriptomic responses to ascorbate deficiency. However, both studies consistently observed an increase in the expression of pathogen-response-related genes, suggesting this may be a core feature of ascorbate deficiency. Since the transcriptomic consequences of ascorbate deficiency remain incompletely understood, further comparative studies under standardized conditions will be important to clarify its context-dependent effects.

Taken together, these findings indicate that the loss of VTC3 has minimal impact on VTC2/GGP, ascorbate redox cycling, or the transcriptome. Nevertheless, the impaired light-induced ascorbate accumulation observed in *vtc3-3* mutants suggests that VTC3 may regulate ascorbate biosynthesis at steps other than VTC2/GGP. One plausible possibility is that VTC3 influences enzymes upstream of VTC2 in the D-Man/L-Gal pathway, such as PMI, PMM, GMP, and GME. These enzymes are responsible for synthesizing GDP-L-galactose, the direct substrate of VTC2/GGP, and reduced activity in any of these steps could limit VTC2 function despite unaltered transcript levels and enzyme activity. Alternatively, VTC3 may affect the supply of essential cofactors required for the optimal activity of VTC2/GGP or its upstream enzymes. For example, VTC2/GGP activity is dependent on the availability of inorganic phosphate. Further studies will be required to identify the precise molecular targets of VTC3 and to clarify how it contributes to the light-responsive regulation of ascorbate biosynthesis. Given that *VTC3* encodes a putative kinase/phosphatase, it may also play broader regulatory roles beyond ascorbate biosynthesis.

Consistent with previous reports (Terai *et al*., 2020; Hamada *et al*., 2023), *vtc2-4* mutants retained the ability to increase ascorbate accumulation in response to light stress (**Figure 1**). We initially hypothesized that this ability might be due to the activation of residual VTC5. However, *VTC5* transcription showed little to no increase in *vtc2-4* mutants compared to the wild type, and GGP activity was nearly undetectable in *vtc2-4* mutants even under high-light stress (**Figure 2**). This could be due to GGP activity in *vtc2-4* being below the detection limit. Indeed, the simultaneous loss of VTC2 and VTC5 results in ascorbate levels below the detection limit, indicating that VTC5 functions as an active enzyme in ascorbate biosynthesis (Dowdle *et al*., 2007; Lim *et al*., 2016). Alternatively, because we used GDP-D-glucose instead of GDP-L-galactose (the physiological substrate, which is commercially unavailable) to measure GGP activity, our assay may not fully reflect endogenous GGP activity. Nevertheless, it is intriguing that ascorbate levels in *vtc2* mutants can increase significantly without a measurable increase in GGP activity. If alternative biosynthetic pathways are not considered, this suggests a mechanism that enhances the GGP response independent of measurable GGP activity. The recently reported metabolon formation of ascorbate biosynthetic enzymes may be involved (Fenech *et al*., 2021), but further experimental validation is required.

Another key objective of this study was to examine the impact of the simultaneous loss of VTC2 and VTC3 on ascorbate pool size. As expected, *vtc2-4 vtc3-3* double mutants exhibited the lowest ascorbate levels reported among viable ascorbate-deficient mutants, retaining only about 14% of wild-type levels (**Figure 4**). Nevertheless, under low-light conditions, *vtc2-4 vtc3-3* double mutants displayed a healthy phenotype similar to wild-type plants. The original *vtc2-1* mutant, isolated via an ethyl methanesulfonate (EMS) mutagenesis screen, has been reported to exhibit dwarfism under standard growth conditions, whereas the T-DNA insertion mutant *vtc2-4*, despite having comparable ascorbate levels, does not exhibit dwarfism (Lim *et al*., 2016). It has been suggested that the dwarfism observed in *vtc2-1* may be due to a mutation in another gene. The relationship between ascorbate levels and growth must be carefully evaluated, as it is highly influenced by growth light intensity. For instance, when grown under relatively higher light conditions (200 µmol photons m^-2^ s^-1^), both *vtc2-1* and *vtc2-4* exhibit similar degrees of dwarfism (Plumb *et al*., 2018). Considering this complexity, our data suggest that under artificial low-light conditions, as little as 14% of wild-type ascorbate levels are sufficient for normal growth in Arabidopsis. This finding provides the idea that the primary reason plants accumulate high concentrations of ascorbate is to protect against oxidative stress rather than to fulfill its coenzyme functions.

The loss of VTC3 significantly impaired high-light-induced ascorbate accumulation in both wild-type and *vtc2-4* backgrounds (**Figure 6**). *vtc2-4 vtc3-3* double mutants maintained extremely low ascorbate levels even under high-light conditions, leading to more severe photooxidative damage than either single mutant (**Figure 5**). In terms of photobleaching severity, *vtc2-4* was more sensitive to high light than *vtc3-3*, and the sensitivity of the double mutant was entirely consistent with its steady-state ascorbate pool size (**Figure 5**). NPQ values were statistically significantly lower in all *vtc* mutants compared to wild-type plants under low-light conditions, with *vtc2-4* and the double mutant exhibiting greater reductions than *vtc3-3* (**Figure 7**). However, no significant difference was observed between *vtc2-4* and the double mutant under low-light conditions, suggesting that NPQ suppression due to ascorbate deficiency may reach a threshold at ∼30% of wild-type ascorbate levels. Interestingly, this difference became more pronounced after high-light exposure, paralleling the fact that *vtc2-4* mutants still exhibited light-induced ascorbate accumulation, whereas this response was severely impaired in the double mutants (**Figure 6**). So far, *vtc1* and *vtc4* mutants have also been used to study ascorbate deficiency, but these mutants have pleiotropic effects unrelated to ascorbate metabolism (Torabinejad *et al*., 2009; Barth *et al*., 2010). In contrast, *vtc2-4* and *vtc3-3* mutants provide a more targeted approach. Since the *vtc2-4 vtc3-3* double mutant retains the characteristics of both single mutants, comparative analysis of these mutants represents a powerful tool for elucidating the physiological roles of ascorbate.

In conclusion, our findings indicate that VTC3 is not required for the expression or activity of VTC2/GGP, ascorbate redox cycling (including ascorbate recycling), or the regulation of the transcriptome. These negative results provide important insight by excluding several plausible roles of VTC3 and thereby advancing our understanding of its physiological function. While the fundamental function of VTC3 remains unresolved, our findings suggest that VTC3 regulates ascorbate biosynthesis at a step other than VTC2/GGP, and future research will focus on validating this notion. Furthermore, this study revealed that the simultaneous knockout of VTC2 and VTC3 exacerbates ascorbate deficiency and increases sensitivity to light stress, likely due to the combined loss of ascorbate biosynthesis capacity (via VTC2) and regulatory modulation (via VTC3), both of which are essential for maintaining sufficient ascorbate levels under photooxidative stress. The newly generated *vtc2-4 vtc3-3* double mutant retains only 14% of wild-type ascorbate levels. Given its distinct physiological characteristics, such as impaired non-photochemical quenching and enhanced photooxidative stress sensitivity, this double mutant, in combination with the single mutants, provides a valuable genetic tool to further dissect the relationship between ascorbate levels and physiological responses. Notably, it represents the most severely ascorbate-deficient viable Arabidopsis mutant characterized to date, potentially enabling the discovery of previously unrecognized roles of ascorbate in plant physiology.

## Supplementary Data

**Supplementary Figure S1** Expression stability of the reference gene *Actin2* across different genotypes and treatment conditions

**Supplementary Figure S2** Expression of *VTC2* and *VTC3* in *vtc* mutant lines

**Supplementary Figure S3** Ascorbate treatment alleviates light stress-induced cell death in the *vtc2- 4 vtc3-3* double mutant

**Supplementary Table S1.** List of primers used

**Supplementary Table S2.** List of genes whose expression is altered in *vtc* mutants and gene ontology enrichment analysis

## Author Contributions

YT, TM, and TI: conceptualization; YT, TM, and TI: methodology; YT, and TM: formal analysis; YT, TM, KA, HN, AH, and TI: investigation; YT, and TM: data curation; YT, and TM: writing - original draft; KA, HN, AH, and TI: writing - review & editing; TM: visualization; TM, TI: supervision; TM, TI: funding acquisition

## Data availability

The data underlying this article are available in the article and its online supplementary material. The raw and processed data of RNA-seq have been deposited in the Gene Expression Omnibus under accession number GSE293768.

## Acknowledgments

We would like to thank Hiromi Ueno, Reona Tanimura, and Natsumi Ito for their technical assistance.

## Abbreviations

APX: ascorbate peroxidase
ASC: ascorbate
DHA: dehydroascorbate
DHAR: dehydroascorbate reductase
GGP: GDP-L-galactose phosphorylase
GME: GDP-D-mannose-3’,5’- epimerase
GMP: GDP-D-mannose pyrophosphorylase
GPP: L-galactose-1-phosphate phosphatase
GSH: reduced glutathione
L-GalDH: L-galactose dehydrogenase
L-GalLDH: L-galactono-1,4-lactone dehydrogenase
MDHA: monodehydroascorbate
MDAR: monodehydroascorbate reductase
NPQ: non-photochemical quenching
PMI: phosphomannose isomerase
PMM: phosphomannomutase
PP2C: protein phosphatase 2C
ROS: reactive oxygen species
uORF: upstream open reading frame
VTC: vitamin c defective

## References

Aarabi F, Ghigi A, Wijesingha Ahchige M, Bulut M, Geigenberger P, Neuhaus HE, Sampathkumar A, Alseekh S, Fernie AR. 2023. Genome-Wide Association Study unveils ascorbate regulation by PAS/LOV PROTEIN during high light acclimation. Plant Physiology doi: 10.1093/plphys/kiad323.

Asada K. 1999. THE WATER-WATER CYCLE IN CHLOROPLASTS: Scavenging of active oxygens and dissipation of excess photons. Annual Review of Plant Physiology and Plant molecular biology 50, 601–639.

Baker NR. 2008. Chlorophyll fluorescence: a probe of photosynthesis in vivo. Annual Review of Plant Biology 59, 89–113.

Barth C, Gouzd ZA, Steele HP, Imperio RM. 2010. A mutation in GDP-mannose pyrophosphorylase causes conditional hypersensitivity to ammonium, resulting in Arabidopsis root growth inhibition, altered ammonium metabolism, and hormone homeostasis. Journal of experimental botany 61, 379–394.

Bournonville C, Mori K, Deslous P, et al. 2023. Blue light promotes ascorbate synthesis by deactivating the PAS/LOV photoreceptor that inhibits GDP-L-galactose phosphorylase. The Plant Cell 35, 2615–2634.

Bulley SM, Rassam M, Hoser D, Otto W, Schünemann N, Wright M, MacRae E, Gleave A, Laing W. 2009. Gene expression studies in kiwifruit and gene over-expression in Arabidopsis indicates that GDP-L-galactose guanyltransferase is a major control point of vitamin C biosynthesis. Journal of Experimental Botany 60, 765–778.

Bulley S, Wright M, Rommens C, et al. 2012. Enhancing ascorbate in fruits and tubers through over-expression of the L-galactose pathway gene GDP-L-galactose phosphorylase. Plant Biotechnology Journal 10, 390–397.

Conklin PL, DePaolo D, Wintle B, Schatz C, Buckenmeyer G. 2013. Identification of Arabidopsis VTC3 as a putative and unique dual function protein kinase::protein phosphatase involved in the regulation of the ascorbic acid pool in plants. Journal of Experimental Botany 64, 2793–2804.

Conklin PL, Gatzek S, Wheeler GL, Dowdle J, Raymond MJ, Rolinski S, Isupov M, Littlechild JA, Smirnoff N. 2006. *Arabidopsis thaliana VTC4* encodes L-galactose-1-P phosphatase, a plant ascorbic acid biosynthetic enzyme. The Journal of Biological Chemistry 281, 15662–15670.

Conklin PL, Norris SR, Wheeler GL, Williams EH, Smirnoff N, Last RL. 1999. Genetic evidence for the role of GDP-mannose in plant ascorbic acid (vitamin C) biosynthesis. Proceedings of the National Academy of Sciences of the United States of America 96, 4198–4203.

Conklin PL, Saracco SA, Norris SR, Last RL. 2000. Identification of ascorbic acid-deficient *Arabidopsis thaliana* mutants. Genetics 154, 847–856.

Dowdle J, Ishikawa T, Gatzek S, Rolinski S, Smirnoff N. 2007. Two genes in *Arabidopsis thaliana* encoding GDP-L-galactose phosphorylase are required for ascorbate biosynthesis and seedling viability. The Plant Journal 52, 673–689.

Fenech M, Amorim-Silva V, Esteban Del Valle A, Arnaud D, Ruiz-Lopez N, Castillo AG, Smirnoff N, Botella MA. 2021. The role of GDP-l-galactose phosphorylase in the control of ascorbate biosynthesis. Plant Physiology 185, 1574–1594.

Foyer CH, Halliwell B. 1977. Purification and properties of dehydroascorbate reductase from spinach leaves. Phytochemistry 16, 1347–1350.

Hamada T, Maruta T. 2024. Measurements of ascorbate and dehydroascorbate in plants using high-performance liquid chromatography. Methods in Molecular Biology (Clifton, N.J.) 2798, 131–139.

Hamada A, Tanaka Y, Ishikawa T, Maruta T. 2023. Chloroplast dehydroascorbate reductase and glutathione cooperatively determine the capacity for ascorbate accumulation under photooxidative stress conditions. The Plant Journal 114, 68–82.

Hossain MA, Asada K. 1985. Monodehydroascorbate reductase from cucumber is a flavin adenine dinucleotide enzyme. The Journal of Biological Chemistry 260, 12920–12926.

Ishida T, Tanaka Y, Maruta T, Ishikawa T. 2024. The D-mannose/L-galactose pathway plays a predominant role in ascorbate biosynthesis in the liverwort *Marchantia polymorpha* but is not regulated by light and oxidative stress. The Plant Journal 117, 805–817.

Iwagami T, Ogawa T, Ishikawa T, Maruta T. 2022. Activation of ascorbate metabolism by nitrogen starvation and its physiological impacts in Arabidopsis thaliana. Bioscience, Biotechnology, and Biochemistry 86, 476–489.

Jander G, Norris SR, Rounsley SD, Bush DF, Levin IM, Last RL. 2002. Arabidopsis map-based cloning in the post-genome era. Plant Physiology 129, 440–450.

Kameoka T, Okayasu T, Kikuraku K, Ogawa T, Sawa Y, Yamamoto H, Ishikawa T, Maruta T. 2021. Cooperation of chloroplast ascorbate peroxidases and proton gradient regulation 5 is critical for protecting Arabidopsis plants from photo-oxidative stress. The Plant Journal 107, 876–892.

Laing WA, Martínez-Sánchez M, Wright MA, et al. 2015. An upstream open reading frame is essential for feedback regulation of ascorbate biosynthesis in Arabidopsis. The Plant Cell 27, 772–786.

Laing WA, Wright MA, Cooney J, Bulley SM. 2007. The missing step of the L-galactose pathway of ascorbate biosynthesis in plants, an L-galactose guanyltransferase, increases leaf ascorbate content. Proceedings of the National Academy of Sciences of the United States of America 104, 9534–9539.

Lim B, Smirnoff N, Cobbett CS, Golz JF. 2016. Ascorbate-deficient *vtc2* mutants in Arabidopsis do not exhibit decreased growth. Frontiers in Plant Science 7, 1025.

Linster CL, Gomez TA, Christensen KC, Adler LN, Young BD, Brenner C, Clarke SG. 2007. Arabidopsis *VTC2* encodes a GDP-L-galactose phosphorylase, the last unknown enzyme in the Smirnoff-Wheeler pathway to ascorbic acid in plants. The Journal of Biological Chemistry 282, 18879–18885.

Lorence A, Chevone BI, Mendes P, Nessler CL. 2004. Myo-inositol oxygenase offers a possible entry point into plant ascorbate biosynthesis. Plant Physiology 134, 1200–1205.

Maruta T. 2022. How does light facilitate vitamin C biosynthesis in leaves? Bioscience, Biotechnology, and Biochemistry 86, 1173–1182.

Maruta T, Sawa Y, Shigeoka S, Ishikawa T. 2016. Diversity and evolution of ascorbate peroxidase functions in chloroplasts: More than just a classical antioxidant enzyme? Plant & Cell Physiology 57, 1377–1386.

Maruta T, Tanaka Y, Yamamoto K, Ishida T, Hamada A, Ishikawa T. 2024. Evolutionary insights into strategy shifts for the safe and effective accumulation of ascorbate in plants. Journal of Experimental Botany 75, 2664–2681.

Mittler R, Zandalinas SI, Fichman Y, Van Breusegem F. 2022. Reactive oxygen species signalling in plant stress responses. Nature Reviews. Molecular Cell Biology 23, 663–679.

Müller-Moulé P, Golan T, Niyogi KK. 2004. Ascorbate-deficient mutants of Arabidopsis grow in high light despite chronic photooxidative stress. Plant Physiology 134, 1163–1172.

Noctor G, Mhamdi A, Foyer CH. 2016. Oxidative stress and antioxidative systems: recipes for successful data collection and interpretation. Plant, Cell & Environment 39, 1140–1160.

Ogura Y, Komatsu A, Zikihara K, Nanjo T, Tokutomi S, Wada M, Kiyosue T. 2008. Blue light diminishes interaction of PAS/LOV proteins, putative blue light receptors in *Arabidopsis thaliana*, with their interacting partners. Journal of Plant Research 121, 97–105.

Pastori GM, Kiddle G, Antoniw J, Bernard S, Veljovic-Jovanovic S, Verrier PJ, Noctor G, Foyer CH. 2003. Leaf vitamin C contents modulate plant defense transcripts and regulate genes that control development through hormone signaling. The Plant Cell 15, 939–951.

Plumb W, Townsend AJ, Rasool B, Alomrani S, Razak N, Karpinska B, Ruban AV, Foyer CH. 2018. Ascorbate-mediated regulation of growth, photoprotection, and photoinhibition in *Arabidopsis thaliana*. Journal of Experimental Botany 69, 2823–2835.

Rahantaniaina M-S, Li S, Chatel-Innocenti G, Tuzet A, Issakidis-Bourguet E, Mhamdi A, Noctor G. 2017. Cytosolic and chloroplastic DHARs cooperate in oxidative stress-driven activation of the salicylic acid pathway. Plant Physiology 174, 956–971.

Shigeoka S, Ishikawa T, Tamoi M, Miyagawa Y, Takeda T, Yabuta Y, Yoshimura K. 2002. Regulation and function of ascorbate peroxidase isoenzymes. Journal of Experimental Botany 53, 1305–1319.

Smirnoff N. 2018. Ascorbic acid metabolism and functions: A comparison of plants and mammals. Free Radical Biology and Medicine 122, 116–129.

Sodeyama T, Nishikawa H, Harai K, Takeshima D, Sawa Y, Maruta T, Ishikawa T. 2021. The d-mannose/l-galactose pathway is the dominant ascorbate biosynthetic route in the moss *Physcomitrium patens*. The Plant Journal 107, 1724–1738.

Tanaka M, Takahashi R, Hamada A, Terai Y, Ogawa T, Sawa Y, Ishikawa T, Maruta T. 2021. Distribution and functions of monodehydroascorbate reductases in plants: Comprehensive reverse genetic analysis of *Arabidopsis thaliana* Enzymes. Antioxidants 10.

Terai Y, Ueno H, Ogawa T, Sawa Y, Miyagi A, Kawai-Yamada M, Ishikawa T, Maruta T. 2020. Dehydroascorbate reductases and glutathione set a threshold for high-light-induced ascorbate accumulation. Plant Physiology 183, 112–122.

Tian T, Liu Y, Yan H, You Q, Yi X, Du Z, Xu W, Su Z. 2017. agriGO v2.0: a GO analysis toolkit for the agricultural community, 2017 update. Nucleic Acids Research 45, W122–W129.

Torabinejad J, Donahue JL, Gunesekera BN, Allen-Daniels MJ, Gillaspy GE. 2009. VTC4 is a bifunctional enzyme that affects myoinositol and ascorbate biosynthesis in plants. Plant Physiology 150, 951–961.

Vidal-Meireles A, Neupert J, Zsigmond L, Rosado-Souza L, Kovács L, Nagy V, Galambos A, Fernie AR, Bock R, Tóth SZ. 2017. Regulation of ascorbate biosynthesis in green algae has evolved to enable rapid stress-induced response via the *VTC2* gene encoding GDP-l-galactose phosphorylase. The New Phytologist 214, 668–681.

Wheeler GL, Jones MA, Smirnoff N. 1998. The biosynthetic pathway of vitamin C in higher plants. Nature 393, 365–369.

Winkler BS, Orselli SM, Rex TS. 1994. The redox couple between glutathione and ascorbic acid: a chemical and physiological perspective. Free Radical Biology & Medicine 17, 333–349.

Yabuta Y, Mieda T, Rapolu M, Nakamura A, Motoki T, Maruta T, Yoshimura K, Ishikawa T, Shigeoka S. 2007. Light regulation of ascorbate biosynthesis is dependent on the photosynthetic electron transport chain but independent of sugars in Arabidopsis. Journal of Experimental Botany 58, 2661–2671.

Yoshimura K, Nakane T, Kume S, Shiomi Y, Maruta T, Ishikawa T, Shigeoka S. 2014. Transient expression analysis revealed the importance of VTC2 expression level in light/dark regulation of ascorbate biosynthesis in Arabidopsis. Bioscience, Biotechnology, and Biochemistry 78, 60–66.

Zechmann B, Stumpe M, Mauch F. 2011. Immunocytochemical determination of the subcellular distribution of ascorbate in plants. Planta 233, 1–12.

