## Supplementary Figures for "Simultaneous knockout of VITAMIN C DEFECTIVE 2 and 3 exacerbates ascorbate deficiency and light stress sensitivity"

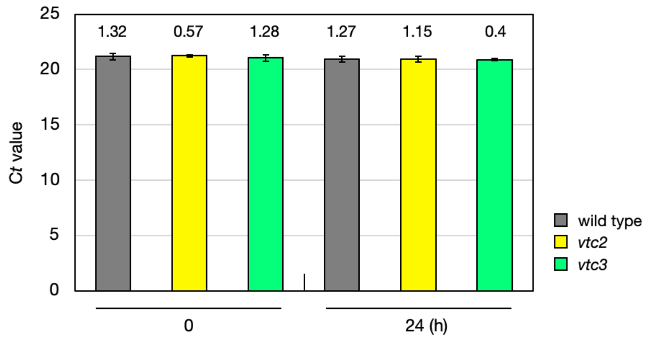


**Supplementary Figure S1** Expression stability of the reference gene *Actin2* across different genotypes and treatment conditions

Bar graphs show the mean C*t* values of *Actin2* across three biological replicates, each calculated from the average of three technical replicates. Error bars represent the standard deviation (SD) of biological replicates. Numerical values above the bars indicate the coefficient of variation (CV, %) calculated for each group. The consistently low CV values (ranging from 0.4% to 1.32%) demonstrate that *Actin2* expression was stable under all experimental conditions, supporting its use as a single reference gene for normalization in this study.

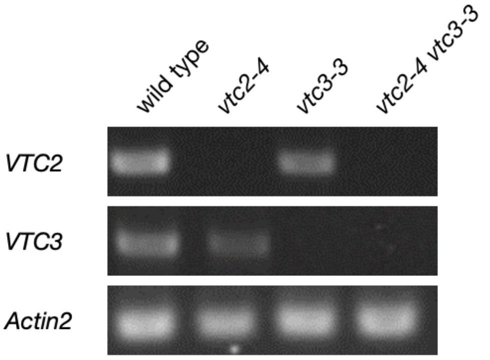


**Supplementary Figure S2** Expression of *VTC2* and *VTC3* in *vtc* mutant lines

Wild-type, *vtc2-4*, *vtc3-3*, and *vtc2-4 vtc3-3* plants were grown on MS plates under low-light (40–60 μmol photons $m^{-2}$ $s^{-1}$) conditions for two weeks. Semi-quantitative RT-PCR data showing expression of *VTC2*, *VTC3*, and *Actin2* (control).

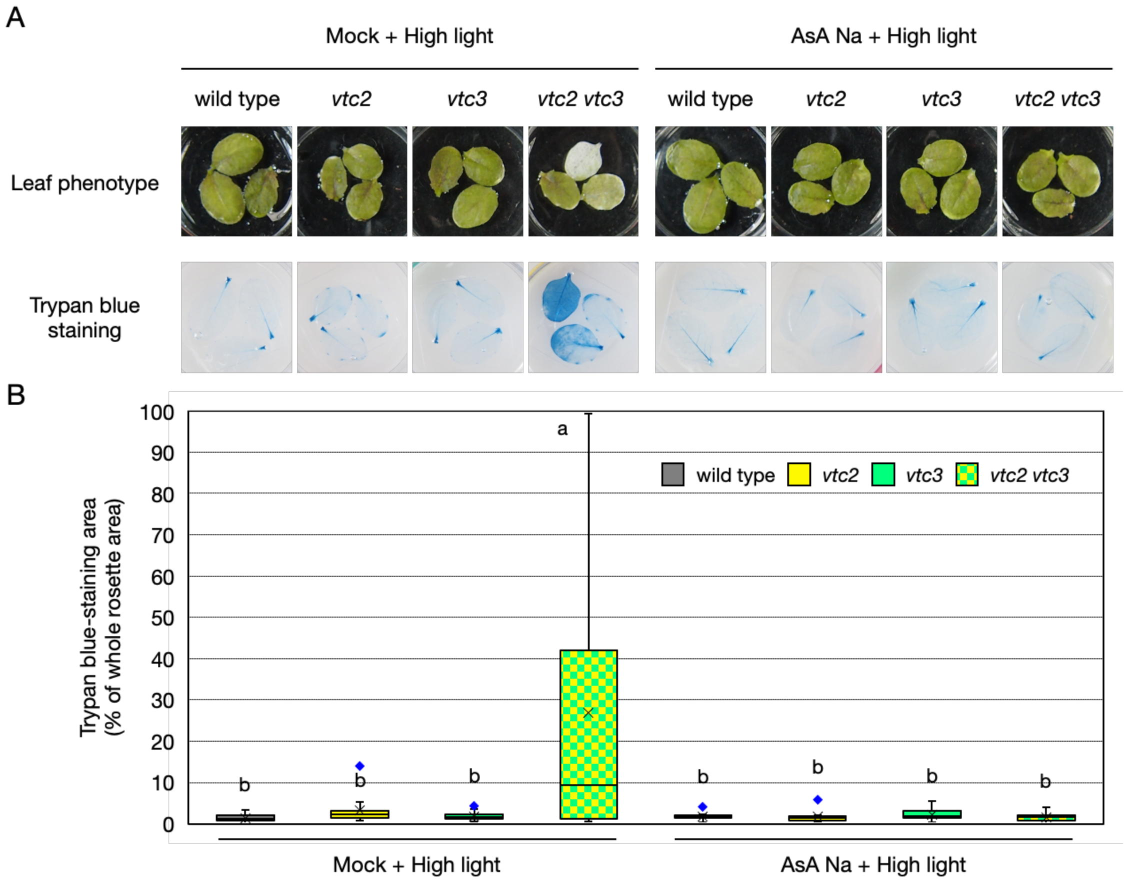


**Supplementary Figure S3** Ascorbate treatment alleviates light stress-induced cell death in the *vtc2-4 vtc3-3* double mutant

Detached leaves from 3-week-old wild-type, *vtc2-4*, *vtc3-3*, and *vtc2-4 vtc3-3* plants were floated on 20 mM MES buffer (pH 5.7) with or without 10 mM sodium ascorbate (AsA Na) and incubated under continuous light for 24 h, followed by high-light stress (1500 µmol photons $m^{-2}$ $s^{-1}$) for 24 h. (A) Top: Representative leaf images after treatment. Bottom: Trypan blue staining to visualize cell death. Clear leaf bleaching and intense trypan blue staining were observed only in *vtc2-4 vtc3-3* leaves without AsA Na, and both were completely suppressed by AsA Na supplementation. (B) Quantification of trypan blue-stained areas using ImageJ software. Box plots show the distribution of data from 15 individual leaves (n = 15), obtained from five biological replicates (three leaves per replicate). Different letters indicate statistically significant differences (Tukey's HSD test, $P < 0.05$).
